# Migration coupled with recombination explains disparate HIV-1 anatomical compartmentalization signals

**DOI:** 10.1101/2023.04.22.537949

**Authors:** Sumantra Sarkar, Ethan Romero-Severson, Thomas Leitner

## Abstract

The evolution of HIV-1 in a host is shaped by many evolutionary forces, including recombination of virus genomes and the potential isolation of viruses into different tissues with compartmentalized evolution. Recombination and compartmentalization have opposite effects on viral diversification, with the former causing global mixing and the latter countering it through spatial segregation of the virus population. Therefore, recombination and compartmentalization together give rise to complex evolutionary dynamics that convolutes their individual effects in standard, bifurcating phylogenetic trees. Although there are various theoretical methods available to infer the presence of recombination or compartmentalization individually, there is little knowledge of their combined effect. To study their interaction and whether that could explain the many disparate results that have been described in the HIV-1 literature, we developed an age-structured forward-time evolutionary model that includes compartments, migration, and recombination. By tracking the evolutionary history of individual virus variants in an Ancestral Recombination Graph (ARG) and resolving the ARG into standard bifurcating trees, we reexamined 771 anatomical tissue pairs infected with HIV-1. Remarkably, we found that recombination can make the resulting bifurcating tree appear more compartmentalized than the virus population actually is. However, we found migration between all 771 tissue pairs, typically at 2.8-4.9×10^−3^ taxa^-1^ day^-1^. Thus, while different point mutations may arise in different parts of the body, migration eventually brings these variants together and recombination merges them into a relatively homogeneous cloud of a universally evolving quasispecies. Modelling this process, we explain the many different results previous research found among distinct anatomical tissues. We also show that the popular Slatkin-Maddison test comes to different results about compartmentalization at the same migration rate depending on the sample size.

**SIGNIFICANCE:** Whether HIV-1 sampled in different anatomical tissues are compartmentalized or not has been a long-standing quest because it relates to both clinical and biological insight. Previous studies of this question have reported disparate results from the same as well as different tissue comparisons. Here, we consolidate and explain these contrasting results with a new evolutionary model that includes compartments, migration, and recombination. We show that no anatomical tissues are fully compartmentalized, and that the migration of HIV-1 between the tissues allows recombination to homogenize the diversity that may arise in separate tissues of an infected person.

## INTRODUCTION

Whether HIV evolution occurs universally throughout the whole human body, or isolated in different anatomical tissues has been a long-standing debate. This debate exists because HIV in the central nervous system (CNS) can lead to neurocognitive impairment [1, 2], in genitalia may be related to sexual transmission [3-5], in any isolated tissue may advance antiviral drug resistance [6], relate to reservoirs that impede cure [7], and may be relevant for future vaccine development.

Naturally, if some parts of the body were truly isolated, then there would be no HIV there, and thus, because we find HIV in all tissues, no part of the body is totally isolated. Nevertheless, some tissues are more separated than others, e.g., the brain is protected by the blood-brain barrier that may restrict HIV from entering the brain from the blood, suggesting some level of restricted gene flow or compartmentalization may occur once some HIV has entered the CNS. Once that has happened, HIV in the CNS would evolve more or less separately, either by genetically adapting to the CNS or simply drifting away neutrally from the ancestral population. Thus, phylogenetic analyses should be able to pick up the signal of compartmentalized evolution by identifying clades that are more or less unique to anatomical tissues. Indeed, there have been a large number of studies that have investigated different tissues in patients. Many found signal for compartmentalization [8-13], and others found no signal for compartmentalization [14-16], even when they studied the same tissue pairs. Sometimes compartmentalization was found in some patients but not others [3, 10, 17]. Yet other studies have shown that there is continuous migration between tissues, implying only a limited degree of or transient localized evolution [11, 18-20].

Complicating things further, Zarate et al. showed that different phylogenetic methods may reach discordant results using the same data, and that topological methods typically were more sensitive in detecting compartmentalization than distance-based methods [21]. Bull et al. showed that even when tests support compartmentalization, collapsing monotypic variants removes the compartmentalization signal and suggests that while some local replication and proliferation may occur, the fact that HIV from different tissues intermingle without tissue-specific mutations still questions compartmentalized evolution [22].

Because all these studies have relied on a standard phylogenetic framework, evolution is assumed to be a bifurcating process, i.e., emerging HIV variants have only one parent. The most popular phylogenetic test that has been applied in HIV compartmentalization research is the Slatkin-Maddison test of restricted gene flow [23, 24]. This, and more modern Bayesian methods, ignores the fact that HIV frequently recombines within the host [25-29]. Critically, recombination and compartmentalization have opposite effects on the overall genetic diversification. Recombination joins separate evolutionary lineages into new chimeric variants while compartmentalization favors isolated, independent accumulation of mutations by spatial segregation of the virus population. Together, recombination and compartmentalization create a complex evolutionary dynamic that convolutes their individual effects in standard, bifurcating phylogenetic trees. Although there are various theoretical methods available to infer the presence of recombination or compartmentalization individually, there is little knowledge of their combined effect.

To study the interaction of recombination and migration between potential anatomical compartments and whether that could explain the many disparate results that have been described in the HIV-1 literature, we developed a forward-time evolutionary model that includes compartments, migration, and recombination. We studied previously collected data representing 61 different human tissues in 771 tissue-pair datasets. We show that recombination, counterintuitively, can make data appear more compartmentalized on a bifurcating tree than it actually is, that popular phylogenetic testing for restricted gene flow is sensitive to sample size, and that there is significant migration between all anatomical tissues in the human body, which leads to near universal evolution of HIV over time.

## MATERIALS AND METHODS

### HIV-1 anatomical tissue sequence data

We collected all previously published HIV-1 datasets that contained ≥2 labelled anatomical tissues and ≥5 HIV-1 sequences per tissue in the LANL HIV database (www.hiv.lanl.gov, collected June 9, 2021), resulting in 842 patient sets covering many different genomic regions (Suppl Fig 1). The LANL HIV database has records from 94 different tissues, but not all of those were represented in patients with ≥2 labelled anatomical tissues. About 87% of the 842 patient sets consisted of only 2 sampled tissues, and the rest had 3-9 sampled tissues. Because some patients’ HIV-1 populations were sequenced in >1 genomic region, we divided the data into separate genomic regions (defined as having a HXB2 genomic start coordinate within 20nt), and filtered further to represent each anatomic tissue and genomic fragment by ≥4 sequences, resulting in 1089 sets with ≥2 labelled anatomical tissues, overall representing 61 different tissues (Suppl Tab 1). To compare the empirical data with the simulated data, we had to restrict to cases where the maximum number of sequences were fewer than 100. This reduced the number of available datasets to 771, still representing 61 different tissues. These sets were used in analyses when the exact length of time since infection was not needed. A subset of 167 tissue pairs also had a known time of infection.

### Simulation model

We developed a forward-time population dynamic model to simulate the change of the effective population size of HIV in a host during chronic infection (Fig 1). Through this model, we investigate the effects of migration and recombination in and between different compartments in an age-structured population. Inspired by experimental observations [30], we assumed that on average older virus variants die earlier in the *i*^*th*^ compartment *N*_*i*_ according to (see derivation in SI)

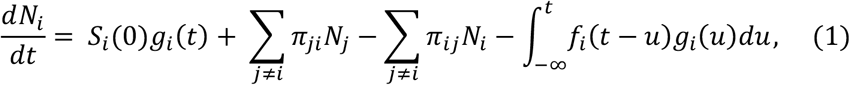

Where *g*_*i*_(*t*) is the time-dependent growth intensity of the population, which we set to be a constant equal to 1 taxon/day. *π*_*ij*_ denotes the per capita migration rate from compartment *i* to *j* measured in units of 1/taxa/day. Finally, *f*_*i*_(*t*) is the extinction age distribution, i.e., the distribution of the time since birth at which a taxon dies and 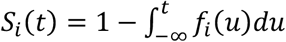 is the survival probability. Motivated by results from HIV-1 neutralization experiments [30], in this paper we assume that the extinction age is Weibull distributed with shape parameter *k* = 3 and scale parameter *λ* = 100.8, corresponding to a mean of 90 days and standard deviation of 33 days.

**Figure 1:**
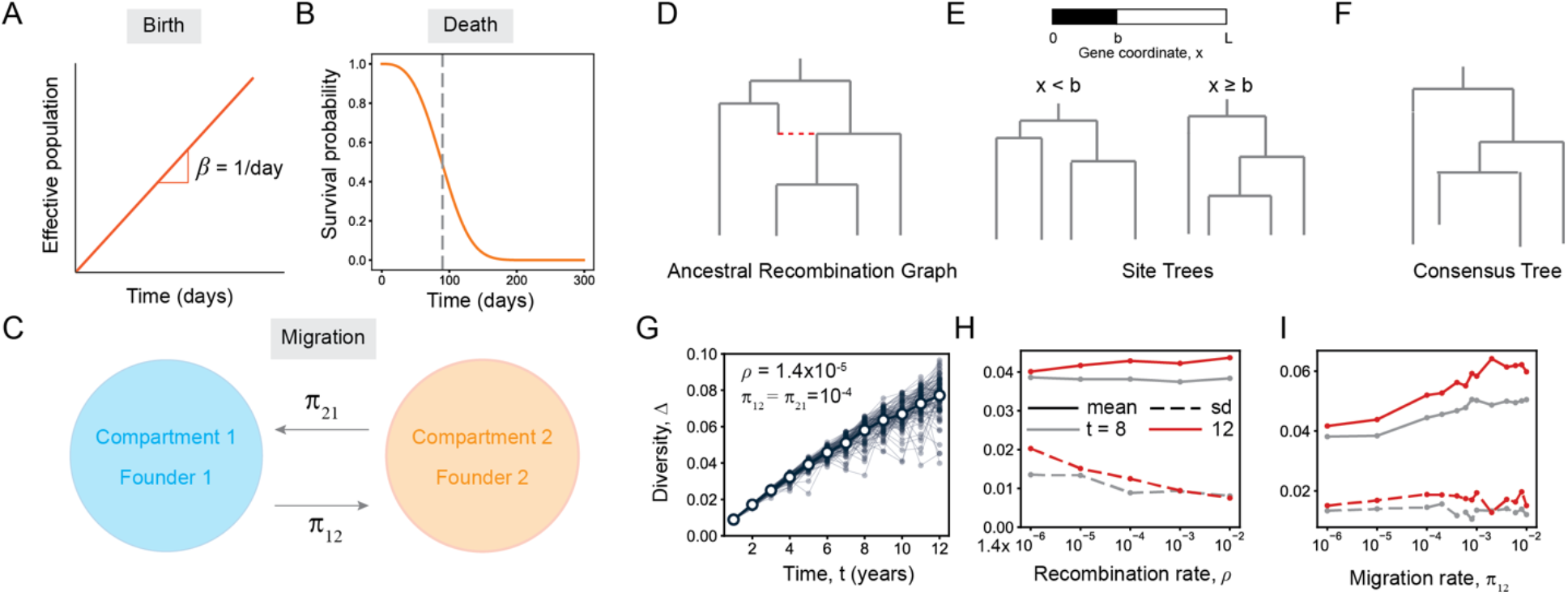
Theoretical model of HIV evolution. (A) Our theoretical framework models the population dynamics of the effective size of an HIV population. The effective population size corresponds to the number of genetically unique HIV-variants in the population. Each individual of this population is a taxon. The effective population size grows linearly at a constant rate of 1/*day*. (B) Due to the interaction with the host immune system, the death of a taxon is age dependent, where older taxa die sooner than younger taxa, as reflected in the survival probability. On average, a taxon dies after 90 days. (C) The migration of taxa between two compartments is described by a random Markovian process, with rates that are identical in both directions (*π*_12_ ≡ *π*_21_). (D) The entire population dynamics can be represented using a network called the Ancestral Recombination Graph (ARG), (E) which can be decomposed into *L* binary site trees on the sampled locus of the viral genome. (F) The consensus binary tree is obtained from a weighted average of the site trees. (G) The genetic diversity, Δ, increases sub-linearly as a function of time. The genetic diversity predicted by the model matches quantitatively with empirical observations. (H) In agreement with Hudson’s model, the mean diversity does not change appreciably with recombination rate *ρ*, and the standard deviation in diversity decreases with increasing recombination rate. (I) The mean diversity increases with increasing migration rate, but the standard deviation does not change appreciably.

Starting from a single identical founding taxon in each compartment, we simulate the model using Gillespie’s exact method [31] for either up to 12 years or, when we compare our model’s predictions to real HIV-1 data, up to the maximum time of sampling in a specific dataset. During the stochastic simulation, the growth, death, and the migration events happen with probabilities proportional to their intensities (Eq. 1). If a growth event occurs, then it can happen either through the splitting of a taxon into two identical taxa or through the non-destructive recombination of two extant taxa into a new taxon, which are again chosen with probabilities proportional to their intensities: *σ*_*i*_*N*_*i*_ for splitting, and 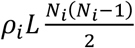 for recombination, where *σ*_*i*_ and *ρ*_*i*_ are the splitting and the recombination rates in the *i*^*th*^ compartment and *L* is the number of nucleotides in the HIV DNA sequence fragment under study. In this paper, focusing on HIV-1 within-host evolution, we set *ρ*_*i*_ = 1.4 × 10^−5^ site^-1^ day^-1^ [25] unless otherwise mentioned and *σ*_*i*_ = 1/day [32]. We explore general theoretical details of this model in a separate study.

### Phylogenetic inference

Our simulation model generates ancestral recombination graphs (ARGs) as a consequence of recombination. While the full simulated ARG is the correct description of the system’s evolutionary history with recombination, ARGs are generally unobservable in practice. Typically, within-host HIV sequences are used to infer a phylogenetic tree to which various tests such as the Slatkin-Maddison test of gene flow [23, 24] are applied. To generate comparable data, we project the ARG a bifurcating tree using the methods described in detail in Castro et al [33]. This involves first decomposing the ARG into a population of bifurcating trees corresponding to each nucleotide site in the aligned sequences, and then computing a consensus tree from the averaged distance matrix over the set of decomposed trees by assuming a minimum evolution criterion, similar to a neighbor-joining tree [34, 35]. When analyzing the real HIV-1 anatomical tissue sequence data, for consistency, we used bionj [36] in the R [37] package ape [38] under a TN93 substitution model, with 1000 bootstrap replicates. While neighbor-joining trees have been shown to be inferior to maximum likelihood trees in HIV research [39], we recently showed that in terms of both topological and branch length summary statistics, they give comparable results when recombination is operating [33].

### Migration rate inference

Our model provides a way to link simple observable quantities such as the minimum number of state changes (i.e. the parsimony score) in an empirical multi-tissue phylogeny to the unobserved true migration rate between those compartments. By running the simulation many times (100 replica for each parameter set) using different migration rates while also simulating the effects of recombination, we obtain the distribution of parsimony scores at each migration rate and obtain the 95% confidence interval, i.e., the range between the 2.5% and 97.5% percentiles. By comparing the model parsimony scores with the bootstrap distribution of the parsimony scores from the empirical data, we can provide an estimate of the range of migration rates consistent with the data. Note, that this approach assumes that the true infection time is known.

To ensure sufficient phylogenetic siqnal, we used two data quality filters. First, that the variance in the distribution of the parsimony score based on 100 bootstrapped phylogenetic trees was less than 90% of the variance in the parsimony score in a set of random trees with the same sampling scheme. If the empirical distribution of the parsimony score computed over bootstrap sampled trees was too variable, we concluded that the phylogenetic signal for the parsimony score was too dependent on a small number of sites. Second, that there was not too much overlap in the empirical distribution of the parsimony score and those generated by a random tree with the same sampling scheme. That is, we concluded that, if the two distributions overlapped by more than 50%, then the data was too close to saturation to get a defensible point estimate of the migration rate. In those cases we could only say that the migration rate must be higher than some number. If a dataset passed both of those tests we included it in our analyses.

## RESULTS

### Recombination reduces the variance but not the mean level of genetic diversity

While recombination can quickly join many polymorphic sites onto a single genome, recombination can only work with the genetic diversity originally introduced by point mutations. Using consensus trees based on site-trees decomposed from ARGs generated by our compartment-recombination model, we found that the mean diversity in a population is essentially unaffected by the recombination rate, while the standard deviation decreases (Fig 1H). These results are consistent with Hudson’s classic coalescent recombination model [40], and suggests that recombination removes mutant outliers of the population, increasing the density of variants more central to the quasispecies, hence reducing the probability of observing a variant with a large set of unique differences stemming for instance from reactivation of latent proviruses [27]. While recombination reduced the variance in diversity by allowing mutations to be easily swapped between different backgrounds, migration instead increased the mean diversity in a compartment without affecting the variance (Fig 1I).

### Recombination makes phylogenetic trees appear more compartmentalized

The degree of compartmentalization for a tissue pair is often assessed based on some interpretation of the parsimony score (i.e., the minimum number of tissue transitions that must have occurred to explain the mixing of labels from different tissues). A low score would imply well-separated compartments, each with unique point mutations typical of a mostly isolated population, while a high score would indicate more migration of genetic variants between the compartments. As expected, we found that the parsimony score increased with elevated migration rates and saturated to a sample size-dependent value at a high enough migration rate (Fig 2).

**Figure 2:**
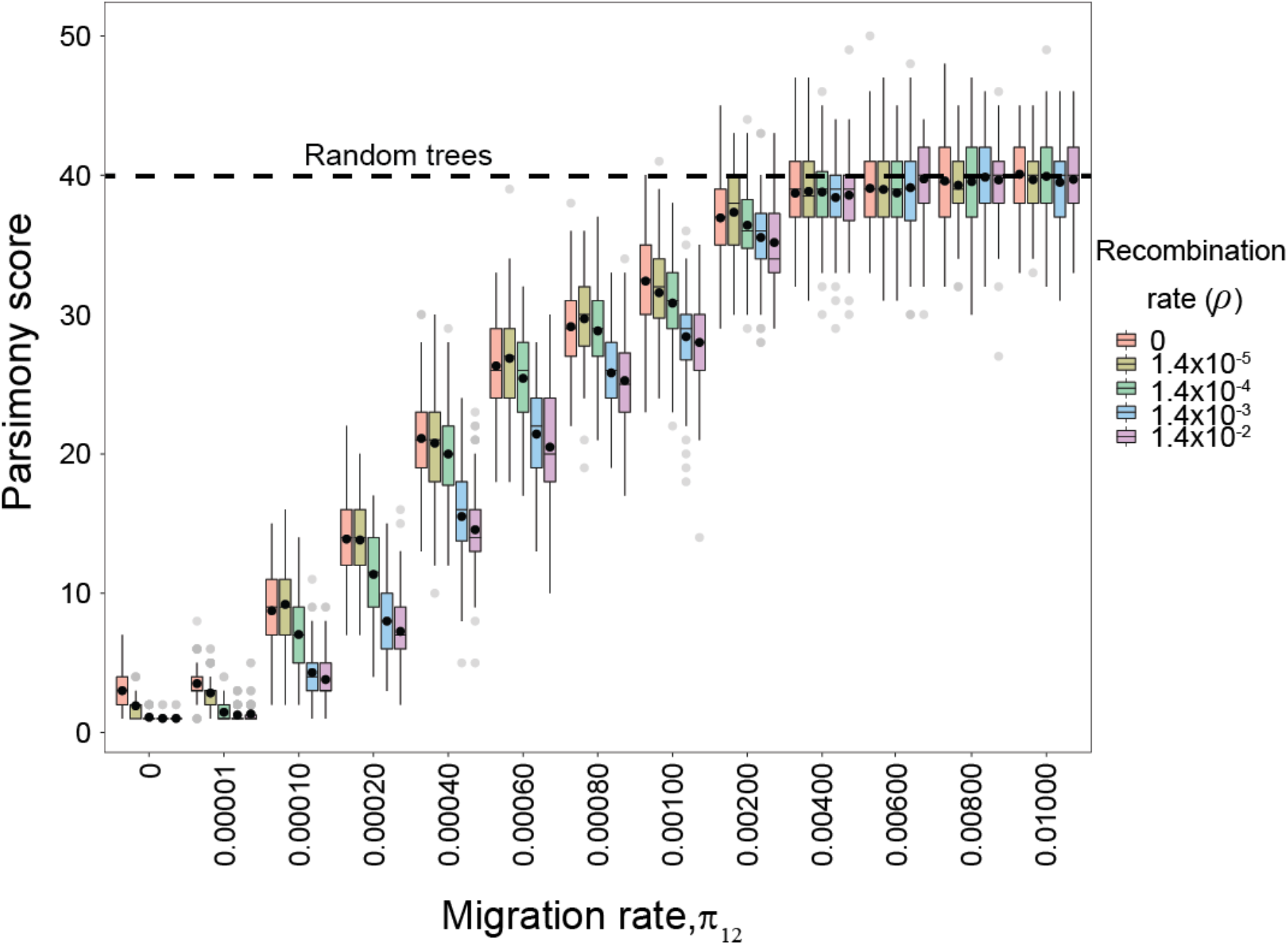
Effects of migration and recombination rates on parsimony score. At all recombination rates, the tissue-label parsimony score increased with increasing migration rates and saturated at a system-dependent maximum value at the mean parsimony score based on random trees given the empirical sequence data size (dashed black horizontal line). At low and intermediate migration rates, the parsimony score depended on the recombination rate: higher recombination rate resulted in a lower parsimony score. For each migration-recombination rate combination, 100 different replicates were analyzed. The horizontal lines in each box shows the median, and the black dot represents the mean parsimony score. Box margins denote the inter-quartile range (IQR), and whiskers ±1.5 IQR. Grey dots indicate values outside the 1.5 IQR range. The origin of nonzero parsimony score at zero migration rate is explained in the SI.

At low and intermediate migration rates, the parsimony score depended on the recombination rate (Fig 2); a higher recombination rate resulted in a lower parsimony score. This makes sense in light of the fact that recombination eliminates outliers (e.g., recent migrants) by integrating their novel mutations into the variants of the receiving population, thereby eliminating the phylogenetic signal of their origin. This effect disappeared at high migration rates as the parsimony score reached that of random trees. Hence, for a given migration rate (that has not reached parsimony score saturation), recombination makes the resulting tree appear more compartmentalized than it actually is, possibly explaining some of the historical difficulty in measuring compartmentalization in HIV-1.

### The parsimony score between anatomic tissue samples predicts migration rate

We can estimate the range of migration rates that are consistent with the tissue-pairs data by finding when our model-based parsimony score overlaps with the parsimony score from bootstrapped empirical data (Fig 3). For a fixed recombination rate, the model-based parsimony score increased sigmoidally as a function of the migration rate (Fig 3A). The possible migration rates for the data were indicated when the 95% confidence intervals of the model-based and the bootstrapped tree distributions overlapped. In Fig 3B, we applied this procedure to estimate the migration rates between HIV-1 infected plasma versus PBMC (peripheral blood mononuclear cells) and plasma versus jejunum (small intestine) in two patients, respectively [41, 42]. With the recombination rate fixed at 1.4 × 10^−5^ site^-1^ day^-1^ [25], as one might have expected, plasma-PBMC showed a higher migration rate as these two tissues are largely intermixed in the body, while plasma-jejunum showed a lower migration rate, consistent with those tissues being more anatomically separated.

**Figure 3:**
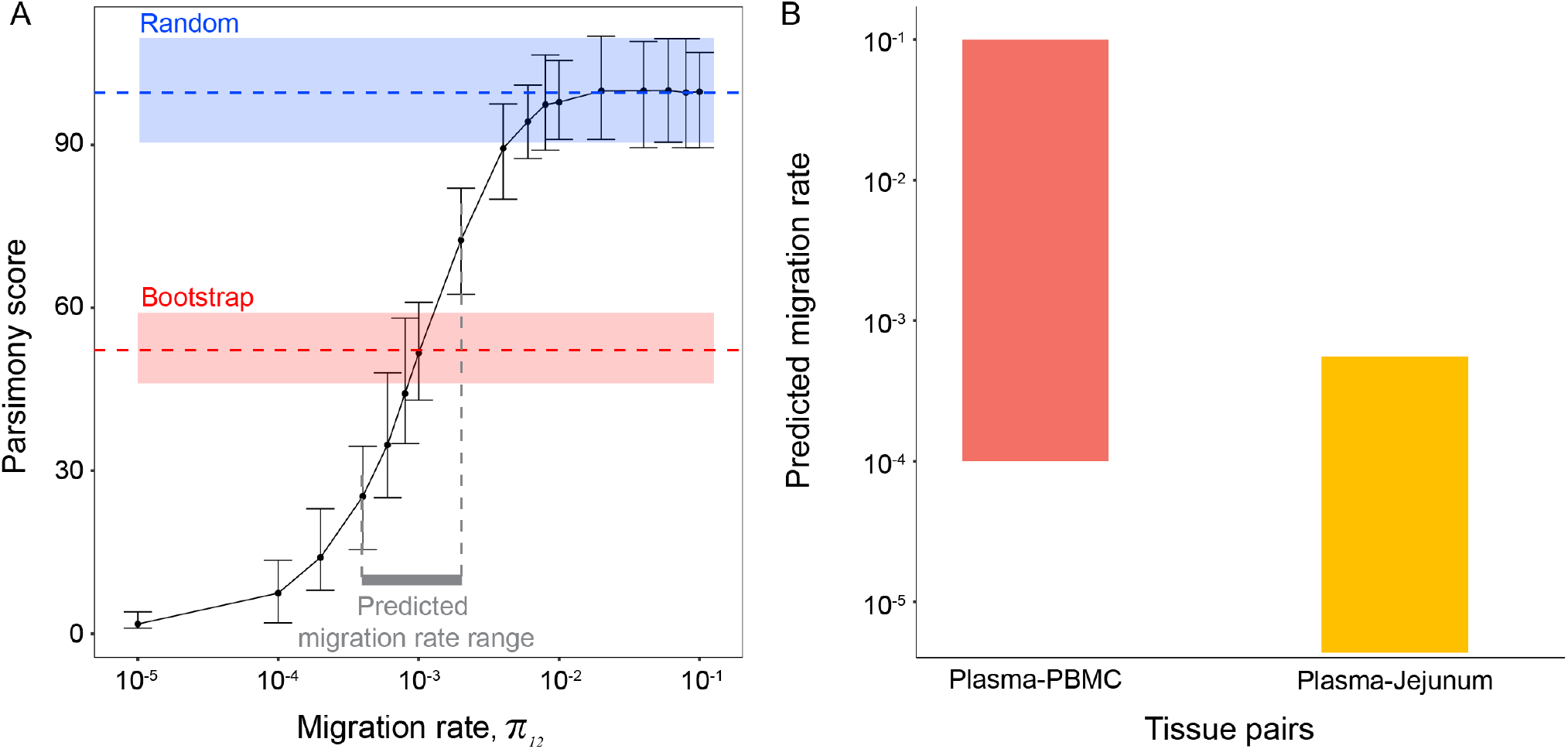
Prediction of migration rates. (**A**) Using our model at a fixed recombination rate, we computed the distribution of the parsimony scores as a function of the migration rate (100 simulations at each migration rate; mean value shown in thin black line with 95% confidence intervals indicated). The red shaded box shows the 95% confidence interval of the bootstrap distribution of parsimony scores from the empirical data, and the blue shaded box shows the 95% confidence interval of the distribution of the parsimony scores of random trees given the empirical data size (labelled Bootstrap and Random). These three distributions are used to estimate the possible migration rates that the data support (see methods for details). The bold red line indicates the possible migration rates supported by the data. (**B**) Applying our method to HIV-1 infected tissue compartments in infected patients; Plasma versus PBMC and Plasma versus Jejunum tissue pairs.

### The compartmentalization signal is affected by sample size

Typically, in studies of compartmentalization, the parsimony score is used in a null hypotheses significance testing framework, like in the classic Slatkin-Maddison test of gene flow [23, 24]. This test computes the probability of an observed parsimony score under the assumption that tip labels are assigned at random (i.e. that there is no “structure” preventing isolates from these distinct tissues from mixing freely as if they came from the same population). If the test is rejected, then some degree of compartmentalization can be inferred. While simple and potentially powerful for some applications, we found that this test is not coherent for the study of HIV dynamics at the tissue scale. Not only was the parsimony score confounded by recombination, but the sample size alone was a strong predictor of whether or not a phylogenetic data set from two distinct tissue types in an HIV-1 infected person was found to be compartmentalized. Analyzing 771 HIV-1 datasets with 4-83 sequences, where two anatomical tissue compartments had been sampled (Fig 4), indeed showed that, for smaller datasets, the p-value distribution was broad, both above and below the typical significance level at 0.05 (Fig 4A). At larger sample sizes most datasets displayed p-values below this level, rejecting the null hypothesis and concluding that some degree of compartmentalization was present. This arises from the fact that the meaning of “compartmentalization” in the Slatkin-Maddison test in terms of the real underlaying migration process is not fixed.

**Figure 4:**
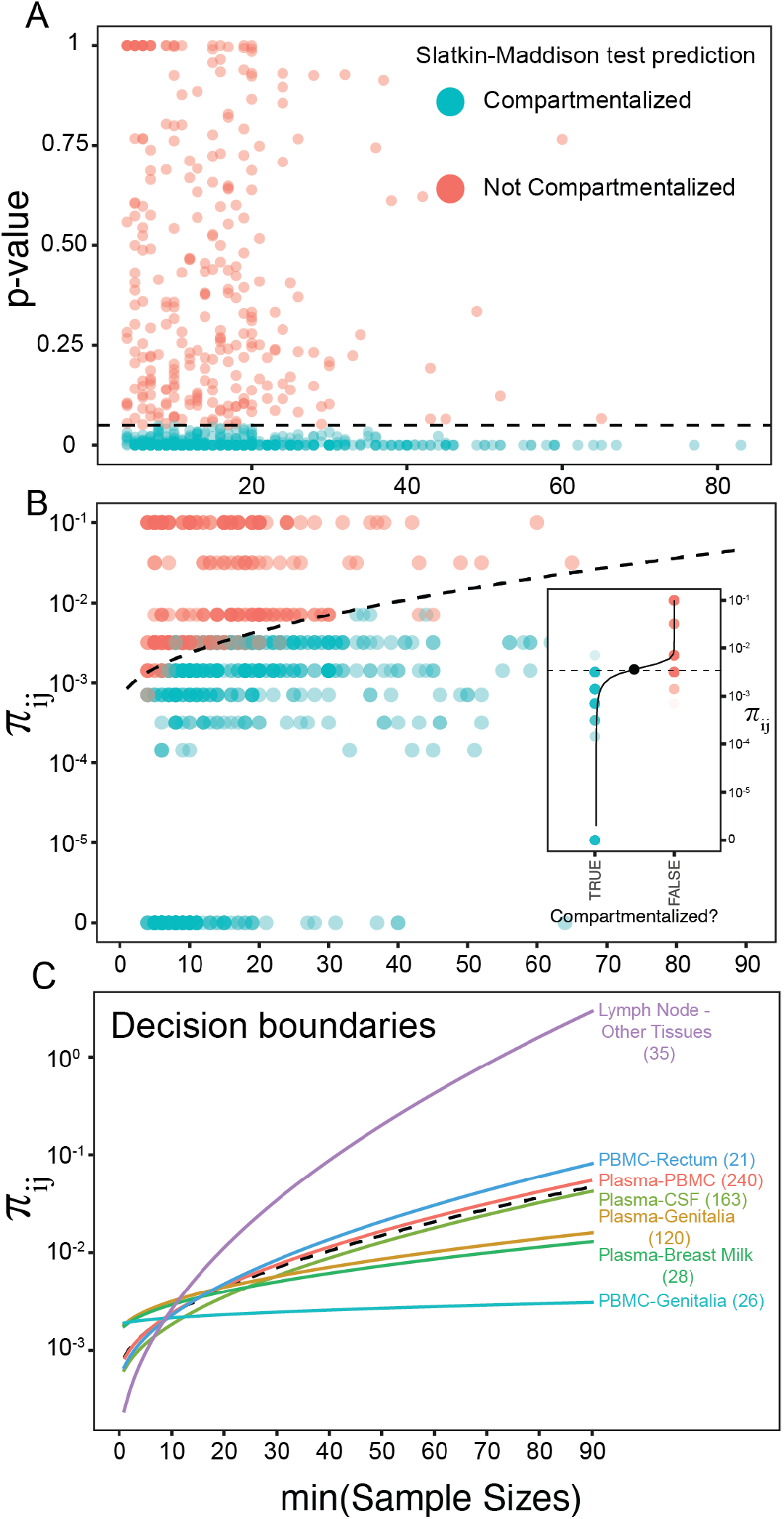
Slatkin-Maddison test results and migration rate inference depends on the empirical sample size. (**A**) Analyzing 771 HIV-1 datasets with two anatomical tissues sampled, the Slatkin-Maddison test showed that the compartmentalization prediction (blue or red dots, see legend) depends on the sample size, i.e., the number of sequences. The dashed line indicates the 0.05 p-value typically used for determining if HIV-1 in the tissues are compartmentalized or not. (**B**) Scatter plot of our model-predicted migration rates versus the sample size, colored by the decision from the Slatkin-Maddison test. A decision boundary, obtained from logistic regression, is indicated as a dashed black line. *Inset:* Scatter-plot of migration rate vs decision when sample size dependence is ignored. A point estimate of the decision boundary (black dot) was obtained using logistic regression (black solid line). (**C**) Decision boundaries estimated from different anatomical tissue pairs (colored lines, number of replicates shown next to tissue labels) compared to the overall decision boundary from panel B. The x-axis shows the size of the smaller sample size of the two compartments compared.

Furthermore, with increased migration rate the parsimony score would saturate at a predicted value for a tree with randomly assigned tip labels, implying that for a given tree, parsimony scores very near the theoretical maximum correspond to as large or larger than some unknown migration rate that will depend on the details of the specific sampling scheme. Thus, two studies using different sampling schemes that both conclude the tip distribution is consistent with random assignment are actually making distinct quantitative claims about the underlying mutation rate.

### No single migration rate differentiates “compartmentalized” from “not compartmentalized”

Crucially, we found that there was no single migration rate that separated “compartmentalized” from “not compartmentalized” in a meaningful way. In the set of 771 paired anatomical datasets we evaluated, the data could be almost perfectly separated into those where the Slatkin-Maddison test was rejected (“no compartmentalization”) or not (“compartmentalization”) based only on the sample size and a certain migration rate (Fig 4B). That is, the Slatkin-Maddison test can be thought of as a binary test of a specific migration rate that depends on the sample size; we call the relationship between the sample size and implied migration rate the ‘decision boundary’.

Figure 4C shows how the decision boundary varied by the type of tissues being compared. While plasma versus other anatomical tissues showed similar decision boundaries, typically, non-plasma versus other (non-plasma) tissues showed either much stronger or much less dependence on migration rate and sample size. In particular, while plasma versus genitalia was similar to plasma versus PBMC, PBMC versus genitalia showed a very different decision boundary, with almost no dependence on migration rate and sample size. Lymph node versus other tissues, in the other observed extreme, showed a much stronger dependence on migration rate and sample size than other tissue pairs. While some of this variation may be due to small number of datasets, it may also indicate potential differences in migration rates between different anatomical tissues.

### Small differences in mean HIV-1 migration rates between different tissue pairs

To assess whether the observed distribution of Slatkin-Maddison test scores for a set of paired tissue datasets can be used to estimate an average migration rate between those two tissues, we first looked at the distribution of p-values for the Slatkin-Maddison test in each tissue pair (Fig 5A). While the proportion of p-values less than the standard 95% level ranged from 0.45 to 0.81, taking the number of datasets into account showed mostly no significant differences (pair-wise, two-sided, two-sample Wilcoxon tests with Bonferroni correction, at *p* = 0.0024, with *α* = 0.05 and *m* = 21 tests). The pairs that showed some difference involved 3 genitalia pairs (p<0.0005) and 1 rectum pair (p=0.001) inconsistently to all other tissues. Interestingly, while plasma-CSF (cerebrospinal fluid) has been claimed to show clear compartmentalization of HIV-1 variants in many previous studies, e.g., [9, 10, 13], this tissue pair did not stand out as clearly compartmentalized in our meta-analysis.

**Figure 5:**
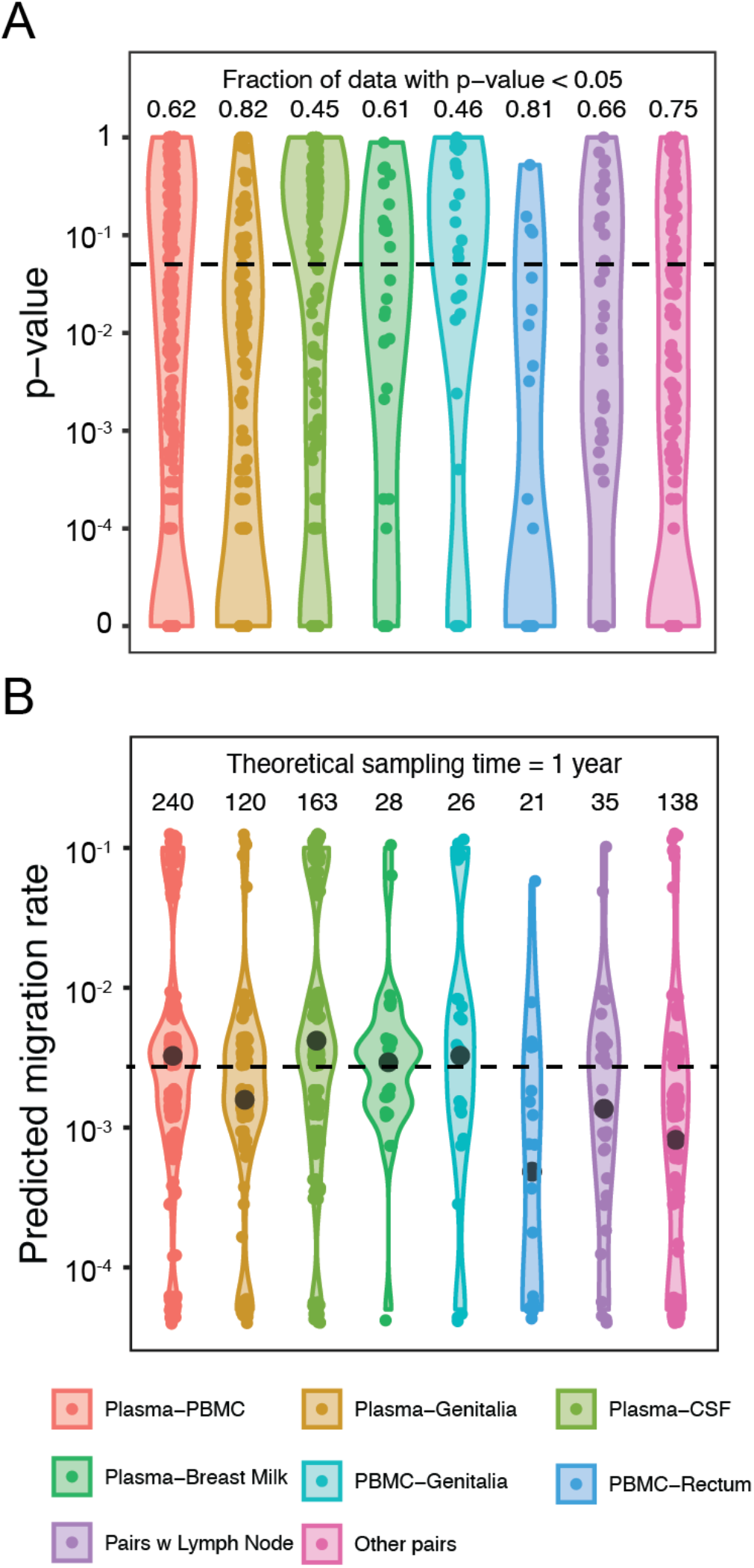
Compartmentalization in tissue pairs. (A) The distribution of p-values obtained from the Slatkin-Maddison test for different tissue pairs (legend). The black dashed line shows the decision boundary at a p-value of 0.05. (B) The estimated migration rates for the same tissue pairs. The mean migration rate is shown with a black dot. The dashed line shows the average point estimate of the decision boundary obtained in 4B. The numbers above the violin plots show the number of datasets available for each tissue pair.

For each tissue pair we estimate a migration rate that was most consistent with the data, as described in the methods. Because we did not know the true infection time in all datasets, we considered simulations where the data were assumed to have been collected 1 or 5 years after infection. The distribution of estimated rates for each tissue pair dataset are shown in figure 5B and the mean of the estimated rates is given for each tissue pair type in table 1. For the most common tissue pair type (Plasma-PBMC, n=240), assuming the sampling took place at 5 versus 1 year from infection reduced the estimated migration rate by 30% leading us to conclude that, in the range of estimated rates that we observe the estimated rates were mostly robust to a wide range of infection times. All else being equal, an individual who has been infected for longer will have a lower migration rate for the same parsimony score.

**Table 1:** The decision point (maximum migration rate) for compartmentalization in different tissue pairs.

| **Tissue Pair** | **Sampling time [years]** | **Number of datasets** | **Decision point (cf. Fig 4B inset)**<br>**[$\times 10^{-3}$ taxa<sup>-1</sup> day<sup>-1</sup>]** |
| --- | --- | --- | --- |
| Plasma-PBMC | 1 | 240 | 3.7 |
| Plasma-PBMC | 5 | 240 | 2.6 |
| Plasma-CSF | 1 | 163 | 3.0 |
| Plasma-Genitalia | 1 | 120 | 4.9 |
| PBMC-Genitalia | 1 | 26 | 3.1 |
| Lymph node-Others | 1 | 35 | 2.8 |
| Plasma-Breast milk | 1 | 28 | 3.2 |
| PBMC-Rectum | 1 | 21 | 3.2 |

Table footnote: The sampling time is at the time since infection in simulations where the number of taxa is equal to the empirical number of sequences from each tissue. The recombination rate was fixed at $1.4 \times 10^{-5}$ site<sup>-1</sup> day<sup>-1</sup> [25].

We found that the mean migration rate for all tissues was 1-7×10^−3^ taxa^-1^ day^-1^, with similar rates across most examined tissue pairs (pair-wise, two-sided, two-sample Wilcoxon tests with Bonferroni correction, *p* = *α*/*m*, with *α* = 0.05 and *m* = 21 tests). The range of possible migration rates we investigated were between 10^−1^ and 10^−5^, which corresponded to situations where compartment labels in the trees would appear completely randomized or nearly completely compartmentalized, respectively. Note that some datasets in the tissue pair collections ended up at these extreme levels (Fig 5B), possibly indicating substantial heterogeneity at the individual host level or heterogeneity in unmeasured confounders such as the distribution of infection times between tissue pair types. The PBMC-rectum tissue pair had a substantially lower estimated mean migration rate at 5×10^−4^ taxa^-1^ day^-1^, but this was represented by the smallest number of datasets (*n* = 21). Finally, we estimated the maximum migration rate that would still give the impression of compartmentalized evolution according to the Slatkin-Maddison test (Table 1). This also indicated relatively small differences (1 to 1.46-fold) among tissue pairs in the order of 10^−3^ taxa^-1^ day^-1^. Thus, overall, while the range of possible migration rates was large for each tissue pair, only relatively small differences in mean migration rates appear to exist between different anatomical tissues in a human with an established HIV-1 infection.

## DISCUSSION

We developed a forward-time within-host evolutionary model that includes potential compartments, migration, and recombination to revisit whether HIV-1 tissues may evolutionarily isolate HIV-1 or not. We show that “compartmentalized” or “not compartmentalized” is not a useful classification, and it unfortunately also depends on the size of the data. Hence, the more sequences that were analyzed, the more likely HIV-1 in one tissue would be found to be “compartmentalized”. Obviously, no tissue is completely compartmentalized, if that was true it would not be possible to harbor (and for us to sample) any HIV-1 at all. Rather, we show that the rate of HIV-1 migration between tissues is what that should be considered. Overall, we found that the migration rate between tissues is on average similar among all tissues we reanalyzed (61 tissues in 771 total pairwise comparisons).

We also show that recombination smooths out diversity extremes, within as well as between tissues. Consequently, recombination homogenizes any population it acts upon. Therefore, counterintuitively, recombination can make tissue specific taxa in a standard bifurcating phylogeny appear more compartmentalized than they actually are. For example, imagine that two tissues each carry unique genetic variants. If a virus from one tissue migrates to the other tissue, and there was no recombination, then the total system would seem less compartmentalized than before the migration. This is the type of event popular compartmentalization detection methods evaluate. Now assume the migrated virus recombines with a variant in the receiving population. If the recombinant off-spring got most of its genetic material from the receiving population variant, then the system would still appear as fully compartmentalized in a bifurcating phylogeny. The same would be true if the recombinant off-spring got most of its genetic material from the donating population variant. With continuous migration and recombination, the uniqueness of the tissue populations would eventually disappear. Thus, while different point mutations may arise in different parts of the body, migration eventually brings variants together and recombination merges these variants into a relatively homogeneous population of an evolving universal quasispecies. Thus, the rate of migration largely determines how soon new variants are integrated into the main cloud of variants.

While previous studies have shown that migration can mix genetic variants sampled from different tissues [11, 18] and replenish tissue compartments [19, 20], to the best of our knowledge, no studies have formally evaluated the combined effects of migration and recombination when evaluating potential compartmentalized HIV-1 evolution. Our model, with compartments, migration, and recombination, allows tracking the evolutionary history of individual virus variants in an Ancestral Recombination Graph (ARG) which is a complicated network of lineage splits (as in a standard phylogeny) and lineage mergers (due to recombination), that adequately describes within-host HIV-1 evolution. While the ARG displays the correct evolutionary history, it is difficult to view, and is still very difficult if not impossible to infer on larger real data [43-46]. On the other hand, standard bifurcating phylogenies cannot properly display recombination, but they are, nevertheless, affected by recombination [33, 47]. Therefore, we used a recently developed method to resolve the ARG into an expected standard bifurcating tree to compare our modelling to the familiar phylogeny [33].

The genetic divergence, which measures the average genetic distance of the extant taxa from the founding taxa, increases linearly with time in our model because the average genetic distance generated by a Poisson distributed molecular clock is always proportional to the temporal distance. The genetic diversity, which measures the average genetic distance in the extant population, changes nonlinearly with time [48]. The nonlinear trend is difficult to surmise from individual trajectories (Fig 1G), which shows large fluctuations in the genetic diversity due to stochasticity from many sources, including population dynamics, sampling, and the measurement of the genetic distance. However, our model predicts that for small time intervals, the average genetic diversity is approximately proportional to the time elapsed, but over longer time intervals the genetic diversity slows down and eventually saturates at a maximum value, depending on the migration rate *π*_*ij*_ and the type of comparison (all compartments, within compartments, between compartments). The average diversity does not depend on the recombination rate (Fig 1H), but the fluctuations decrease with increased recombination rates, which are expected from the Hudson model of recombination [40].

Our modelling system does not include selection, another important factor that can affect the trends of divergence and diversity. While the immune system puts a universal pressure on the virus, affecting variants in all anatomic tissues, it is possible that there are other tissue specific pressures that may encourage different mutations to emerge in different tissues. Previous work showed that lung and blood tissues did not display selection for specific variants [18]. However, compartmentalized neutral evolution would also result in different mutation profiles in different tissues, just at a different rate than by selection. Regardless of how mutations may accumulate in different tissues, migration would eventually bring these variants together, and recombination would mix them into chimeric forms. Thus, while selection could alter the rate at which variants and chimeric forms may emerge, our model shows good qualitative and quantitative agreement of the divergence and diversity trends to experimental observations [48], suggesting that our model captures the HIV evolution in the chronic phase. While the overall trend in genetic diversity of the plasma HIV-1 population is arguably determined by continuous selection of HIV variants during the chronic phase, inspired by observations from neutralization assay experiments [30], our model captures this process in a neutral age-structured population where newer HIV variants are probabilistically more likely to survive than older HIV variants.

In conclusion, we have shown that while different point mutations may arise in different parts of the body, migration eventually brings these variants together and recombination merges them into a relatively homogeneous cloud of a universally evolving quasispecies. The use of standard bifurcating trees to study compartmentalization is misleading because recombination can make the resulting bifurcating tree appear more compartmentalized than the virus population actually is, and we found migration between all 771 tissue pairs in the metadata analysis. Thus, the combined effects of migration and recombination cannot be ignored when studying HIV-1 evolution in different anatomical sites.

## Supplementary information

**Figure S1:**
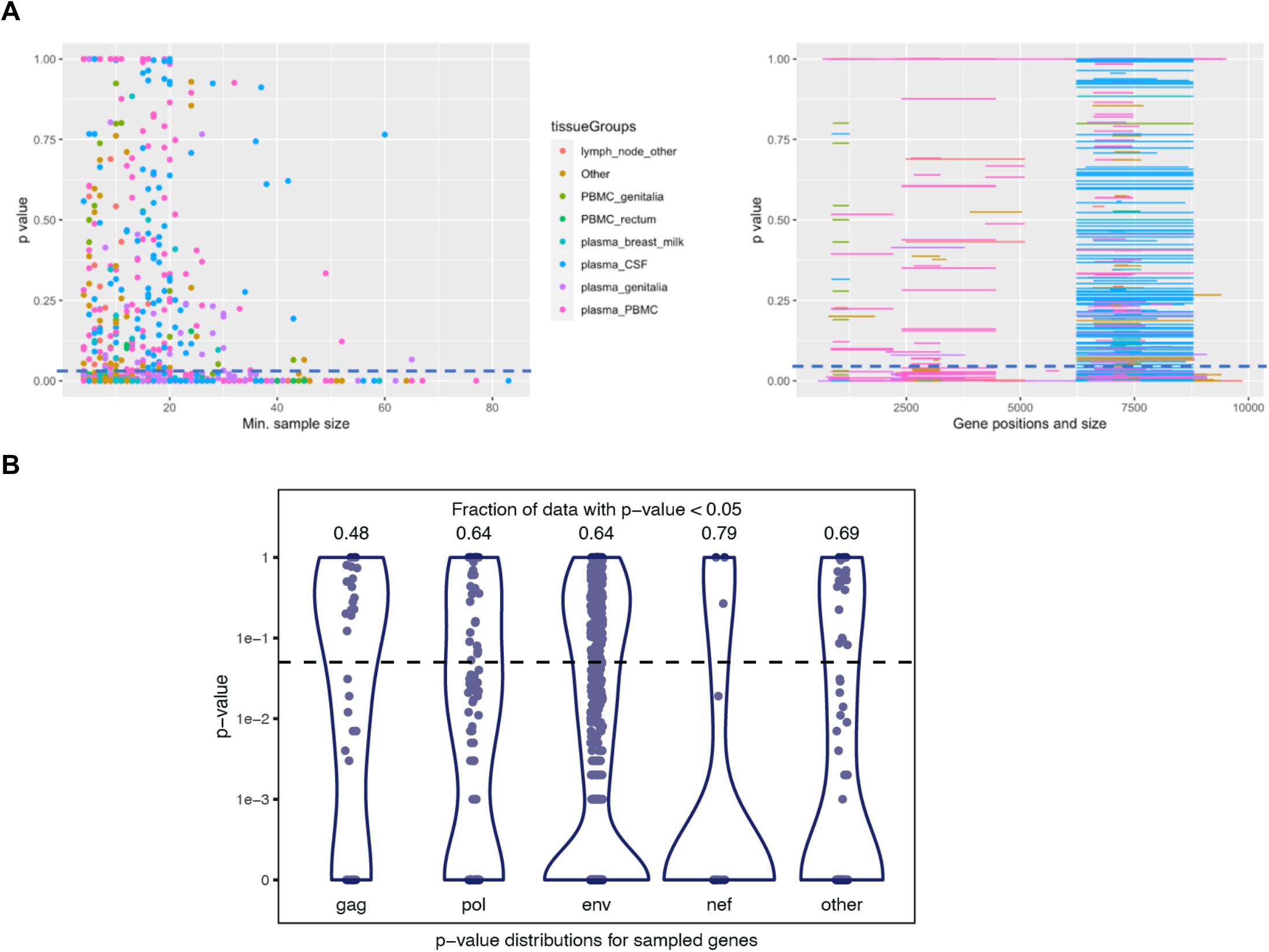
**(A)** (left panel) The p-value from Slatkin-Madisson tests as a function of the tissue with the smaller sample size in tissue pair comparisons. (right panel) The p-value as a function of the sampled gene location. The size of the sampled fragment is shown by the length of the line. P-value lower than 0.05 indicates significant compartmentalization and above this value, no compartmentalization is supported. The detection of compartmentalization depends strongly on the sample size, but shows no dependence on the genetic loci and length. **(B)**The p-value from the Slatkin-Madisson test for various sampled gene location. P-value lower than 0.05 indicates significant compartmentalization and above this value, no compartmentalization is supported. There is no discernible dependence of compartmentalization with gene location.

**Table S1.**
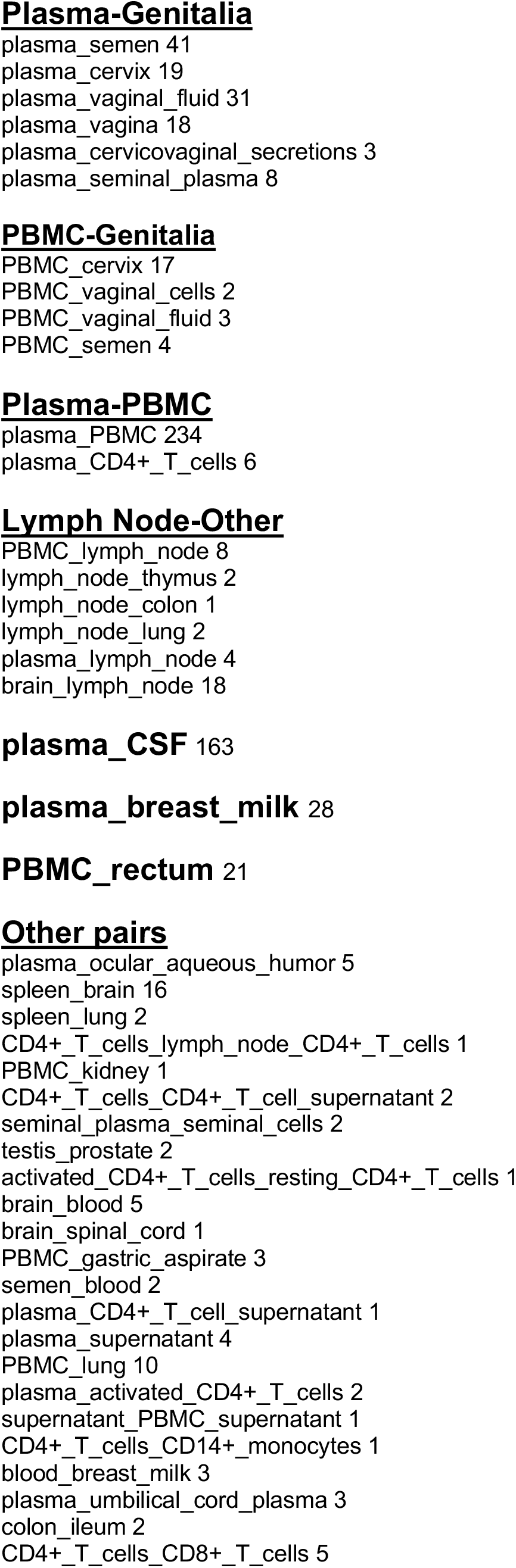
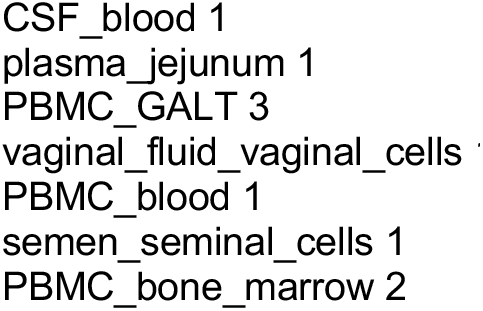
Grouping of tissues.

## Derivation of the equation governing age-dependent population dynamics

The equation governing population dynamics with age-dependent death rate is given by:

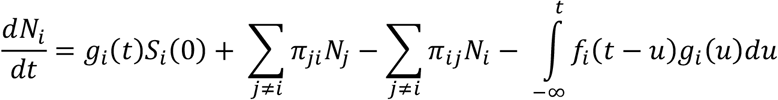

In this equation, *N*_*i*_(t) is the population in the *i*th component; *g*_*i*_ (*t*) is the time dependent growth rate of the taxa in the same component. *π*_*ij*_ denotes the per capita migration rate from compartment *i* to *j* measured in units of 1/taxa/day. Finally, *f*_*i*_(*t*) is the extinction age distribution, i.e., the distribution of the time since birth at which a taxon dies and 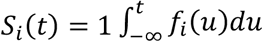 is the survival probability. The birth-death process is nonmarkovian because of the age-dependence, but the migration between compartments is a memoryless markovian process.

To get intuition about the implications of this equation, consider a simplified situation with no migration.

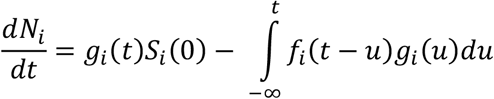

We can make exact calculation in the extreme case when all taxa dies precisely after a certain number of days, say *T*. In this situation, *f*_*i*_ (*t* − *u*) = *δ*(*t* − *u* − *T*). Hence,

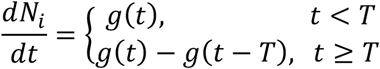

Let’s assume that the virus grows at a constant rate when *t* > 0. That is, *g*(*t*) = *g*Θ(*t*). Integrating the equation for population dynamics we get:

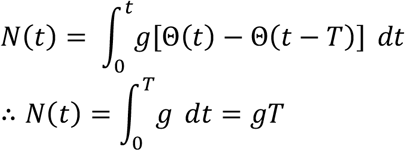

Hence, the total number of taxa remains constant, as expected.

### Derivation of the equation

To derive this equation, we write down the equation governing population at age *a* at time *t, n*(*a, t*). The equation is:

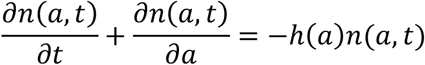

In this equation, *h*(*t*) = *f*(*t*)/*S*(*t*) is the *hazard rate*. To get the equation govering the total population, we integrate this equation over *a*:

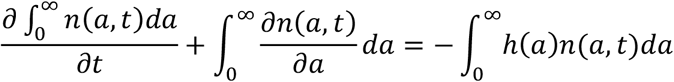

That is:

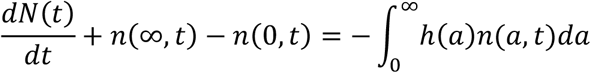

Now, let’s consider the special case of *linear growth*, such that the growth intensity does not depend on the current population and is given by *g*(*t*). Therefore, under this model *n*(*a, t*) = *g*(*t* − *a*)*S*(*a*), i.e., it simply counts how many taxa were born at time *t* − *a*, multiplied by the survival probability after time *a*. Hence, for this special case, corresponding to HIV taxa growth, we can rewrite the above equation in the following simple form.

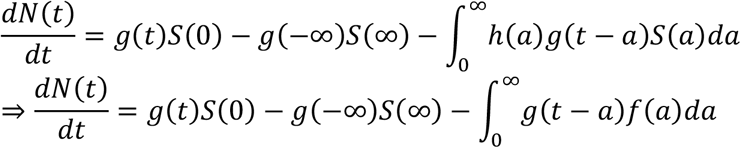

Change variable *u* = *t* − *a* and make the realistic assumption that *S*(∞) = 0. With these changes, we get:

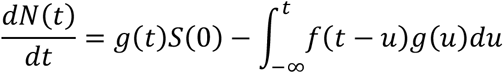

Because the migration rate from compartments *i* to *j* is *π*_*ij*_, the intensity of migration is *π*_*ij*_*N*_*i*_. Therefore, the total change in population in compartment *i* only due to migration is:

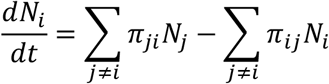

Taken together, we get the equation for population dynamics governing our compartmentalized system.

### Simulation details

We use an agent based model to simulate the population dynamics. We start with two identical founding taxa in two compartments. At the beginning of the simulation, we sample from the survival time distribution the number of days after which these founding taxa die. We put them in a sorted queue where the taxa that is next to die is placed at the front of the queue. This process is repeated as soon as a new taxa is born. This queuing process ensures that the death process is age-dependent. All taxa grow at a rate of *g* =1 taxa/day. The migration rates and the recombination rates are varied from simulation to simulation. For a given iteration, the intensity of the growth and the migration process is calculated. The intensity for the former is *g*, for both compartment 1 and 2 because of the linear growth, and for the latter for compartment *i* is ∑_*j* ≠*i*_ *π*_*ji*_ *N*_*j*_, i.e. *π*_12_*N*_1_ for compartment 1 and *π*_21_*N*_2_ compartment 2. Hence, the total intensity is:

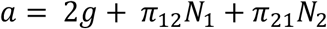

This total intensity is used in the exact Gillespie stochastic algorithm to sample a next event time and the next event. If a growth event is chosen as the next event, splitting or recombination events are chosen based on their intensities, as described in the methods section. The simulation ends at a predetermined maximum time. The simulation box is periodically sampled nondestructively to perform phylogenetic analysis.

## Origin of nonzero parsimony score at low migration rate

We start our simulation with two identical taxa on two compartments. As a result, in the absence of much mixing of the gentic information, such as through recombination, the evolution of the taxa are nearly identical in both compartments, even when we include stochasticity. Hence, when we construct the phylogenetic tree by sampling both compartments, there is sufficient interpenetration of the branches. As a result, the parsimony score is nonzero. At high recombination rates, genetic material within a compartment mix sufficiently well. In the presence of stochasticity, this mixing is sufficient to separate the branches, such that the parsimony score becomes zero, as expected.

When we introduced nonzero genetic distance between the founding taxa, the parsimony score at low migration rate became zero for all recombination rates. Because we chose the genetic distance between the founders arbitrarily and there was no rigorous basis of choosing one genetic distance over the others, we decided to make the parsimonious choice that the founders are genetically identical. This choice does not affect our results in this paper.

## Notes

### Competing Interest Statement

The authors have declared no competing interest.

